# Trade-offs in Robustness to Perturbations of Bacterial Population Controllers

**DOI:** 10.1101/2020.06.04.134932

**Authors:** Cameron McBride, Domitilla Del Vecchio

## Abstract

Synthetic biology applications have the potential to have lasting impact; however, there is considerable difficulty in scaling up engineered genetic circuits. One of the current hurdles is resource sharing, where different circuit components become implicitly coupled through the host cell’s pool of resources, which may destroy circuit function. One potential solution around this problem is to distribute genetic circuit components across multiple cell strains and control the cell population size using a population controller. In these situations, perturbations in the availability of cellular resources, such as due to resource sharing, will affect the performance of the population controller. In this work, we model a genetic population controller implemented by a genetic circuit while considering perturbations in the availability of cellular resources. We analyze how these intracellular perturbations and extracellular disturbances to cell growth affect cell population size. We find that it is not possible to tune the population controller’s gain such that the population density is robust to both extracellular disturbances and perturbations to the pool of available resources.

## 1. Introduction

Many exciting applications exist for engineered biological systems in fields ranging from medicine to the environment [1]–[4]. In order to create sufficiently sophisticated engineered systems that can be applied in practice, substantial research has gone into techniques to scale up the size and complexity of genetic circuits. One major hurdle to scaling up a circuit’s size is resource sharing, where different circuit components become undesirably coupled through the host cell’s pool of resources [5], [6] and affects cell growth [7]–[9]. Solutions to the resource sharing problem have recently appeared. Some solutions make genetic modules robust to resource fluctuations with feedback control [10], [11], some engineer separate pool of resources for the engineered circuits [12], and others distribute the circuit across multiple cell strains [13], [14].

Some highly complex circuits have been created by distributing genetic circuits across different cell strains [14], [16]–[22]. Many of these systems utilize microfluidic platforms or spacial separation to isolate bacterial strains from one another. However, if multiple cell strains are not spatially separated, population control is required to maintain a desired cell population size [20]. In future applications where genetic circuits are distributed across cells and each cell also contains a population controller, the population controller’s robustness to perturbations in cellular resources, for instance, those applied by the other genetic circuits, will influence the overall system’s function.

In particular, in a setup where the population controller operates concurrently with other genetic circuits, such as with a sensor or a more sophisticated circuit that releases pharmaceutical proteins, perturbations to the pool of resources due to these circuits affect the population controller as a disturbance. Meanwhile, changes in the availability of nutrients in the environment, for example, due to the presence of other cell strains competing for nutrients, affect the dynamics of the cell population size that the controller is attempting to regulate as shown in Figure 1. Here, we specifically focus on the potential trade-off between robustness to extracellular perturbations due to the presence of other cells competing for nutrients (*W* in Figure 1), and to intracellular perturbations to the pool of cell’s resources due to the expression of other genetic circuits (X in Figure 1). In related work, [15] considers a decrease in growth rate due to the expression of enzymes and finds a condition on the genetic circuit parameters where dividing the production of a metabolite across multiple cell strains results in higher yield; however, perturbations in the availability of cellular resources and environmental disturbances are not considered.

**Fig. 1.**
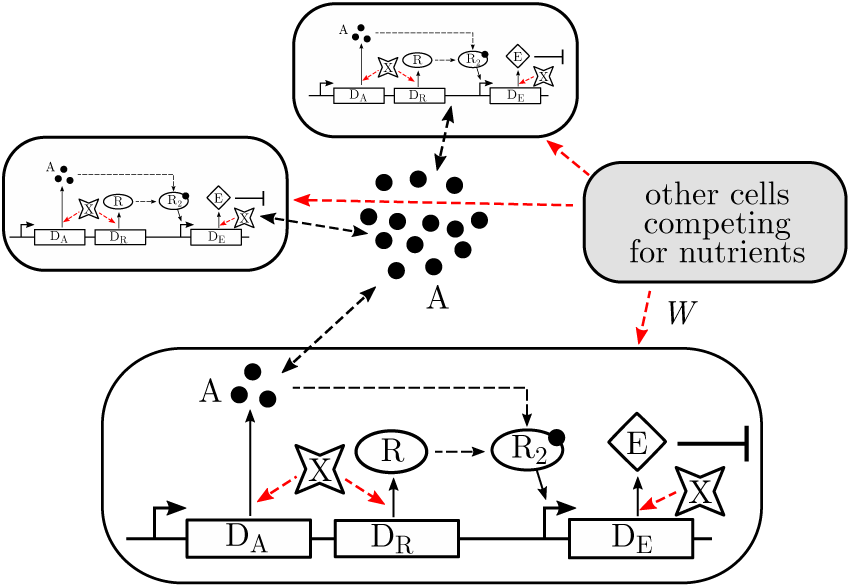
Population control genetic circuit. The metabolite A is produced from DNA D_A_. The metabolite A diffuses through the cell membrane into the environment. A also diffuses into the cell and binds to R to become R_2_. The R_2_ complex then activates the production of the killing gene E, which increases the cell death rate. Each protein production step requires cellular resources X to function. Other cells in the environment compete for nutrients with the strain of interest, resulting in the disturbance *W*. We consider *W* and fluctuations in X as disturbance inputs.

This paper is organized as follows. In Section II, we create a mechanistic model for cell growth and the population controller. In Section III, we present our main results by analyzing this model for its sensitivity to disturbances in the environment and to fluctuations in available cellular resources. We conclude in Section IV.

## II. Model of a Population Control System

We consider the cellular population control circuit of [26], shown in Figure 1. This controller has been shown to maintain a constant number of cells in a bacterial population significantly below the carrying capacity of the environment under controlled laboratory conditions. This population control architecture has been used in most of the applications of genetic circuits distributed across multiple cells [20]. We create a model to describe the population dynamics, then we will model the dynamics of the population controller. We will use this system model to analyze how the number of cells in the population changes with extracellular disturbances, for example, due to competing cell strains, and with intra-cellular disturbances, for example, due to the expression of additional synthetic genes within the cell [5].

Considering the situation shown in Figure 1, let the number of cells in the bacterial population of interest be *N*. We model the population size with the logistic growth model, which is standard for bacteria [26], [27]. Then, the dynamics of the cell population size are given as

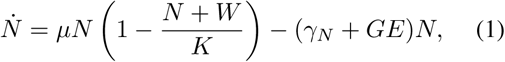

where *µ* is the maximum growth rate of the cell line, *K* is the carrying capacity of the environment, *γ*_*N*_ is the basal rate of cell death, *E* is the concentration of the killing protein E, and *G* is the effect of the concentration of the killing protein on the population death rate. Additionally, *W* represents an environmental disturbance such as the presence of other cell strains, which reduces the available nutrients to our cell strain of interest.

Next, we model the biochemical species that control the cell population size *N* through the production of the killing protein E as shown in Figure 1. In this system, each cell produces the small molecule A at a constant rate. Here, we model the generation of this molecule as a simple gene expression process. Then, molecule A diffuses through the cellular membrane into the surrounding media. We assume that the diffusion of A occurs quickly and the media is well mixed so the concentration of A is uniform throughout (both within and outside the cell). As the concentration of A becomes large due the presence of a large number of cells, A diffuses into cells and binds to R, which is expressed at a constant rate. The active complex of R with A is represented by R_2_. The R_2_ complex then activates the production of the killing protein E, which kills some of the cells, decreasing the number of cells in the population [26]. This population controller structure is based off a common quorum sensing motif found in multiple natural bacteria [24], [25].

In the model, we include resources X required for gene expression as indicated in Figure 1. Specifically, we lump together the processes of transcription and translation [28], so that we can view X as a lumped resource required for protein production. Additionally, we consider the free resources as an input for simplicity. More realistic models may consider the free resources as a state that depends on all proteins being produced in the cell [28] with the activation or repression of other circuits appearing as the new disturbance, imparting a change in X. We ignore any interaction between resource fluctuations and cell growth and assume that protein degradation and dilution remain constant and are independent of population size. Under these assumptions, the chemical reactions describing the biochemical controller are given as

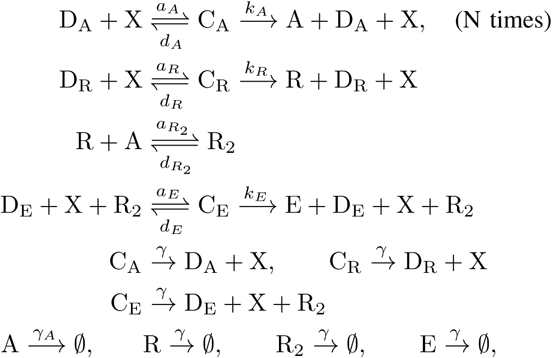

where D_A_, D_R_, and D_E_ are the DNA molecules that produce A, R and E, respectively. Since A diffuses through the cellular membrane and the concentration of A is uniform throughout the media, the production rate of A is proportional to the cell population size *N* since each cell produces A and *N* is dimensionless. Then, using the law of mass-action [29], the dynamics of the concentration of the biochemical controller species are given as

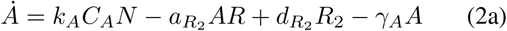

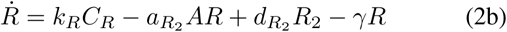

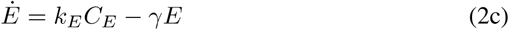

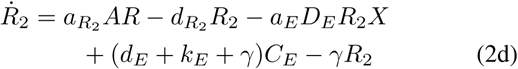

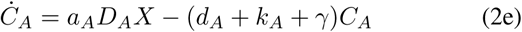

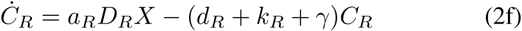

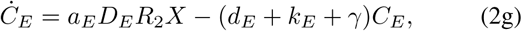

with the conservation laws for the DNA

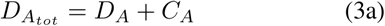

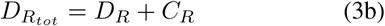

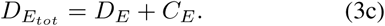

Next, we utilize timescale separation to reduce the dimension of (2), since we are interested in the dynamics of this system on the slow timescale. Since binding and unbinding reactions occur much faster than protein production and degradation [28], i.e. 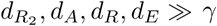, we define a small parameter 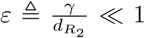 and the parameters 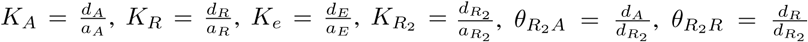, and 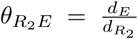. Then *R*_2_, *C*_*A*_, *C*_*R*_, and *C*_*E*_ are fast variables, while *A* and *R* are mixed variables, and *E* is a slow variable. We apply a change of coordinates and define the new stat0065s *z*_*A*_ = *A* + *R*_2_ + *C*_*E*_ and *z*_*R*_ = *R* + *R*_2_ + *C*_*E*_, which are slow variables. Then, the system can be written in standard singular perturbation form [30] as

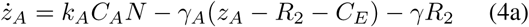

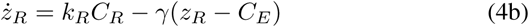

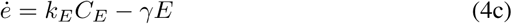

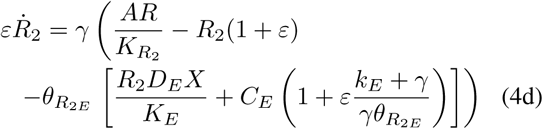

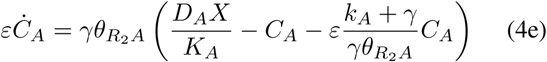

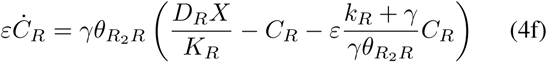

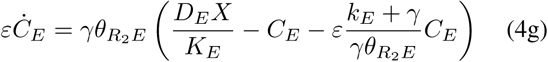

with *z*_*A*_, *z*_*R*_, and *E* as slow variables. Next, we let *ε* → 0. Then, the slow manifold is given as

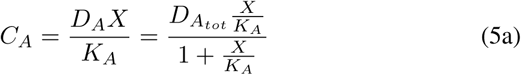

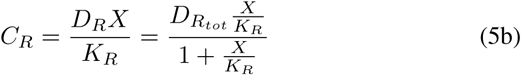

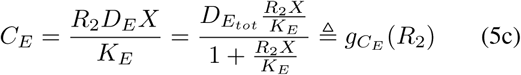

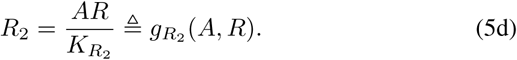

The slow manifold is exponentially stable, which is verified by checking that the eigenvalues of the Jacobian of the fast states evaluated on the slow manifold (4d)–(4g) have uniformly negative real parts [30]. The Jacobian computed on the fast manifold is given as

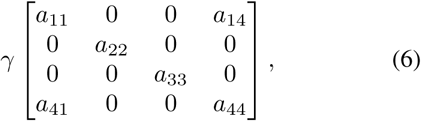

where 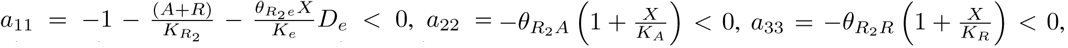 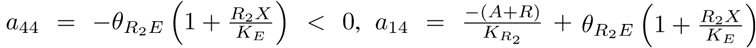, and 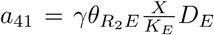. It can be shown that the real part of all eigenvalues of (6) are uniformly negative if *a*_22_, *a*_33_ *<* 0, which are always satisfied since concentrations are always nonnegative, and if *a*_11_*a*_44_ *> a*_14_*a*_41_. The latter inequality reduces to

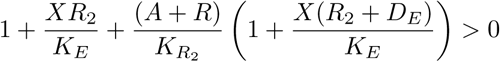

which is also always satisfied since concentrations are nonnegative. Thus, the slow manifold is exponentially stable. Next, substituting (5) into the dynamics of the slow variables with the conservation laws (3), and defining 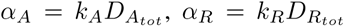, and 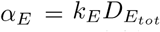, we have the reduced dynamics of the slow variables given by

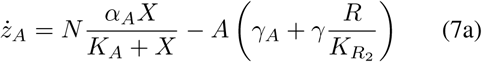

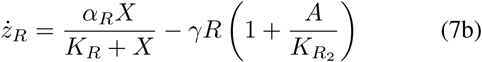

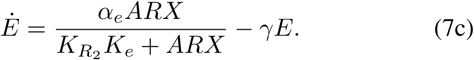

We now change variables back to the original variables *A* and *R* where the relation of the time derivatives between *z*_*A*_ and *z*_*R*_ to *A* and *R* are given as

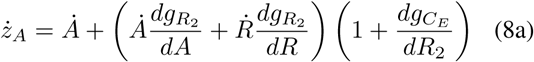

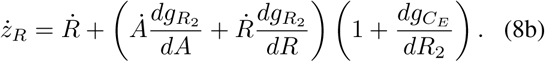

Solving (8) for 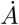 and 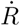, we have

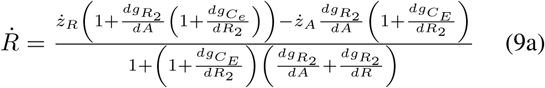

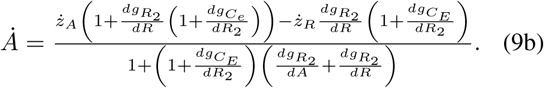

Next, we substitute (7) into (9). We assume that 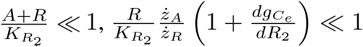, and 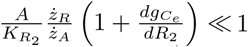, which can be guaranteed if binding between A and R is sufficiently weak. Then, system (9) takes the approximate form

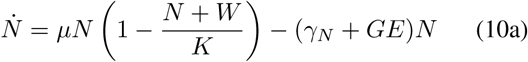

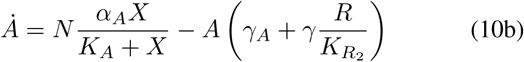

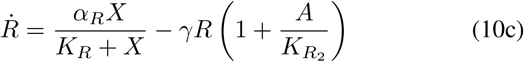

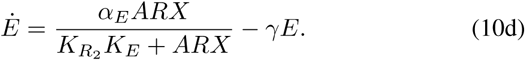

In this model, *N* is the output variable of interest while *W* and *X* are disturbances, which affect the population dynamics and the biomolecular feedback controller, respectively, as shown in Figure 2. In the following section, we analyze the sensitivity of the output *N* of (10) with respect to the disturbances *X* and *W*.

**Fig. 2.**
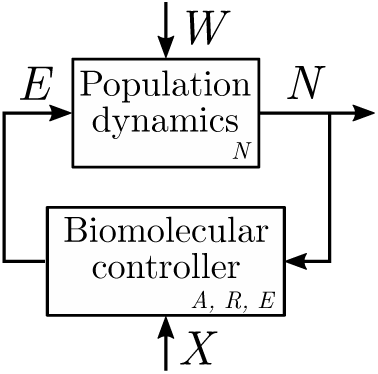
Block diagram of nonlinear system (10) with disturbance inputs *W* and *X* and output *N*. The population dynamics block has state *N*, and the biomolecular controller block has states *A, R*, and *E*.

## III. Closed Loop Sensitivity Analysis

We now analyze the sensitivity of the cell population density *N* with respect to the environmental disturbance *W* and the internal resource disturbance *X*. To achieve this, we linearize the nonlinear system (10) about an equilibrium point and compute the transfer functions from *W* and *X* to *N*. We then use these sensitivity transfer functions to analyze the trade-off between robustness to step disturbances in *W* and in *X*.

### A. Linearization

Let (*N*_0_, *A*_0_, *R*_0_, *E*_0_) be an equilibrium point of system (10) in the positive orthant corresponding to constant nonnegative input levels *X*_0_ and *W*_0_. Let *δ*_*W*_ = *W* − *W*_0_ and *δ*_*X*_ = *X* − *X*_0_ be small perturbations (disturbances) to the nominal inputs *X*_0_ and *W*_0_, and let *δ*_*N*_ = *N* − *N*_0_, *δ*_*A*_ = *A* − *A*_0_, *δ*_*R*_ = *R* − *R*_0_, and *δ*_*E*_ = *E* − *E*_0_ be the corresponding perturbations to the state of system (10). Then, the dynamics of these perturbations can be obtained by the linearization of system (10) about equilibrium point (*N*_0_, *A*_0_, *R*_0_, *E*_0_) as

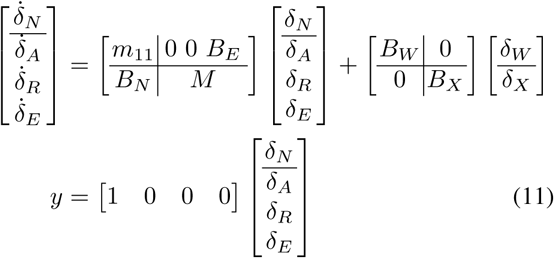

where 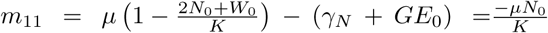, which simplifies using the equilibrium point constraint in (10a). Additionally, *B*_*E*_ = −*GN*_0_, 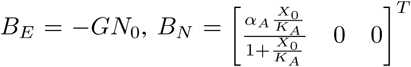, and

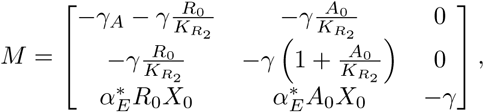

where 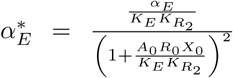. The disturbance input matrices are given as 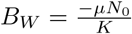, and

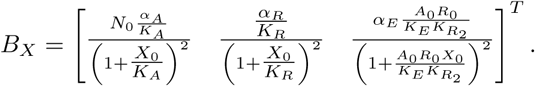

Next, we take advantage of the structure of (11) and rewrite the system as a feedback interconnection between of the growth dynamics *δ*_*N*_ and the biomolecular controller dynamics as in Figure 2. The population controller takes *δ*_*E*_ and *δ*_*W*_ as inputs and gives *δ*_*N*_ as the output, and the biomolecular controller takes *δ*_*N*_ and *δ*_*X*_ as inputs and gives *δ*_*E*_ as the output. Then, we have the state space representation of the population dynamics

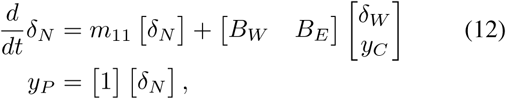

and the state space representation for the biomolecular controller as

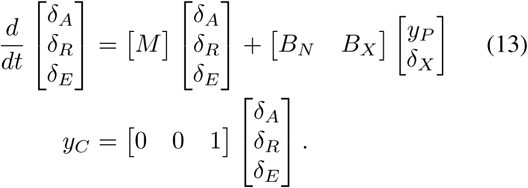

### B. Problem formulation

We wish to analyze how the disturbances *δ*_*W*_ and *δ*_*X*_ affect the output *δ*_*N*_. In this system, it is possible to change the degradation rate *γ*_*A*_ of *A* by changing the pH of the media [26]. Additionally, we will show that 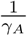 is related to the closed loop gain, and will examine how changing *γ*_*A*_ affects the sensitivity of the output *δ*_*N*_ to disturbances *δ*_*W*_ and *δ*_*X*_. To this end, we compute transfer functions from *δ*_*W*_ and *δ*_*X*_ to *δ*_*N*_ using (12) and (13). Using (12), the transfer functions from *δ*_*E*_ to *δ*_*N*_ and from *δ*_*W*_ to *δ*_*N*_ are given by *P* (*s*) = [1](*s* −*m*_11_)^−1^*B*_*E*_ and *P*_*W*_ (*s*) = [1](*s*− *m*_11_)^−1^*B*_*W*_, respectively, using the standard formula. Then, simplifying, we have

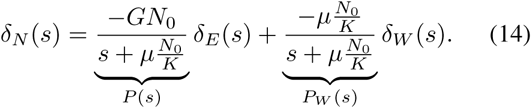

Additionally, the transfer function of the biomolecular controller from *δ*_*N*_ to *δ*_*E*_ using (13) is 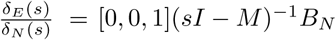, which simplifies to become

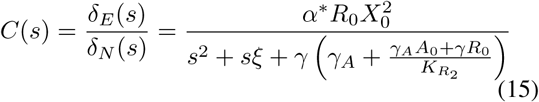

where 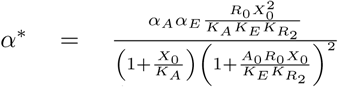 and 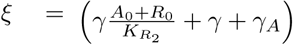. The transfer function from the disturbance *δ*_*X*_ to *δ*_*E*_ is given as 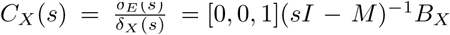. To simplify and for ease of analysis later, we pull *C*_*X*_ (*s*) through *C*(*s*) and define 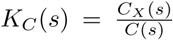, which is shown in Figure 3 and is given by

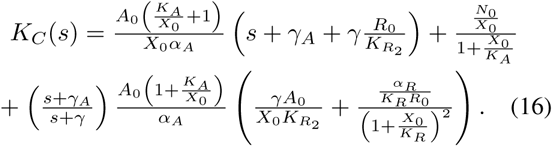

**Fig. 3.**
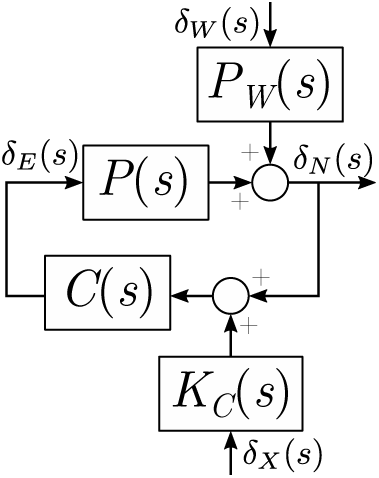
Block diagram of the closed loop system rearranged such that the perturbation *δ*_*X*_ (*s*) appears on the feedback path. This form is helpful for analyzing the closed loop sensitivity of *δ*_*N*_ (*s*) to perturbations in the combination of *δ*_*W*_ (*s*) and *δ*_*X*_ (*s*).

In the closed loop system, we wish to evaluate how tuning the parameter *γ*_*A*_ affects the performance of the biomolecular controller. To this end, we calculate the loop gain of the closed loop system *L*(*s*) = *P* (*s*)*C*(*s*), which is given as

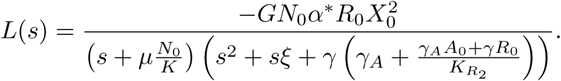

Thus, the loop gain *L*(*s*) may be tuned by varying *γ*_*A*_.

Next, we analyze a weighted sensitivity function that is a linear combination of the sensitivity of *δ*_*N*_ to *δ*_*X*_ and to *δ*_*W*_ and show in Theorem 1 that this function has a strictly positive lower bound.

#### Definition 1.

Let the magnitude of the sensitivity of the output *δ*_*N*_ to the disturbance *δ*_*W*_ be

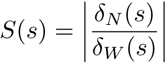

and the normalized magnitude of the sensitivity of the output *δ*_*N*_ to the disturbance *δ* be

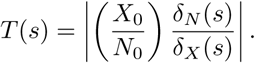

The total weighted sensitivity function is defined as

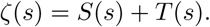

The value of *ζ*(*s*) determines the magnitude of the response of *δ*_*N*_ with respect to the combination of the disturbances *δ*_*W*_ and *δ*_*X*_.

#### Theorem 1.

*Consider system* (11) *where δ*_*W*_ *and δ*_*X*_ *are step disturbances, then the weighted sensitivity function ζ evaluated at s* = 0 *has the lower bound*

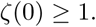

*Proof.* First we evaluate *S*(*s*) at *s* = 0. From the block diagram in Figure 3, 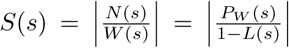. At *s* = 0, *P*_*W*_(0) = −1, so 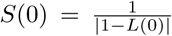. Next, we evaluate *T*(*s*) at *s* = 0. From the block diagram in Figure 3, 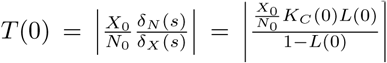 where *K*_*C*_(*s*) is given in (16). To find a bound for *T* (0), we note that 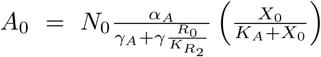 from (10b). Then, substituting and evaluating (16) at *s* = 0 and simplifying, *K*_*C*_(0) is given as

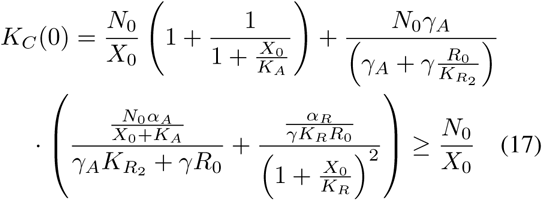

since all concentrations are nonnegative. Substituting the relations for *T* (0), *S*(0), and the inequality (17) into the definition of *ζ* and simplifying, we have

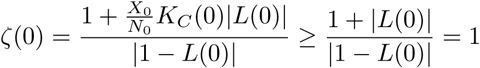

since *L*(0) ≤ 0. □

Theorem 1 shows that for any stable equilibrium point of (10), it is impossible to tune the population controller to reject step disturbances from both the environment *δ*_*W*_ and resource fluctuations *δ*_*X*_, i.e. *S*(0) and *T* (0) cannot both be made small simultaneously. In Figure 4, *S*(0), *T* (0), and *ζ*(0) are shown for different values of the parameter *γ*_*A*_, and *ζ*(0) is bounded from below by 0 dB. Future work may consider the system for *s* ≠ 0 and compare the sensitivities at each value of *s*.

**Fig. 4.**
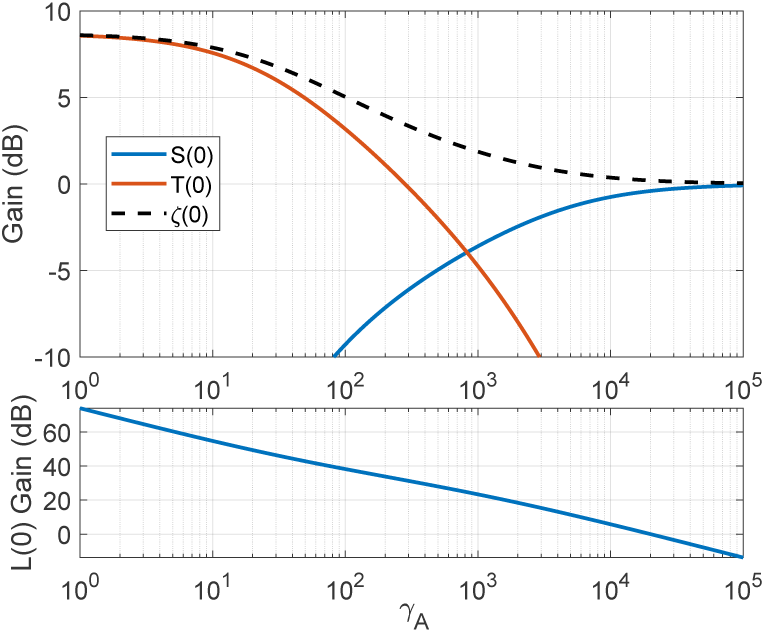
Values of the sensitivity functions *S, T*, and *ζ* about the equilibrium point of (10) evaluated at *s* = 0 for different values of *γ*_*A*_. For these parameters, the magnitude of the loop gain |*L*(0)| decreases as *γ*_*A*_ is increased. *ζ*(0) shows the lower bound of 0 dB derived in Theorem 1. Parameters used for the simulations are 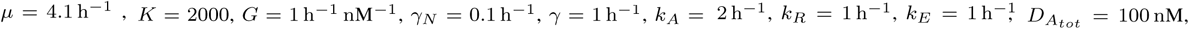 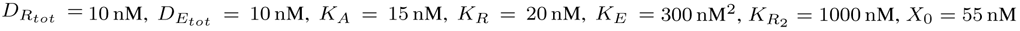. The equilibrium point of the system for all values of *γ*_*A*_ shown exhibit asymptotic stability.

### C. Remarks on Stability of the Closed Loop System

The calculation of sensitivity functions in Section III only have meaning when the closed loop system is stable, since the output after a perturbation must remain within a neighborhood of the equilibrium point. Here we find conditions where the closed loop is guaranteed to be stable using the closed loop linearized transfer function. From (14) and (15), the closed loop characteristic polynomial is given as

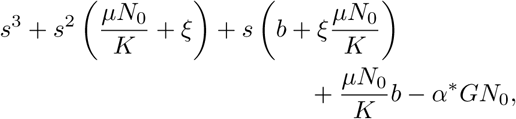

where 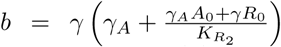. From the Routh-Hurwitz criterion for a 3rd degree polynomial, the closed loop system is stable if and only if

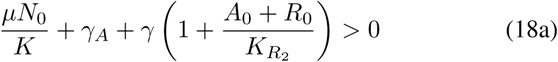

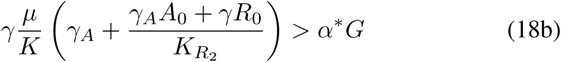

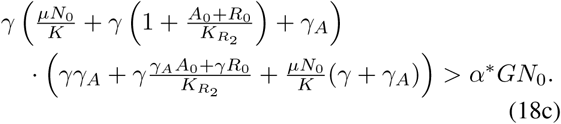

If we also assume that 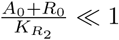 and 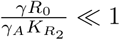, as before, then (18) simplifies to

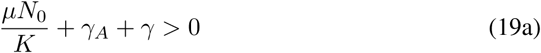

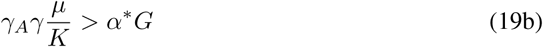

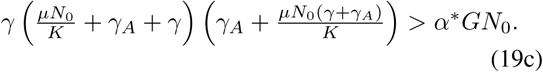

Since 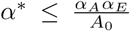 and *N*_0_ ≤ *K*, if 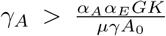 and 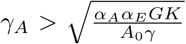, then all inequalities in (19) are satisfied. Since *A*_0_ is bounded away from 0, then the closed loop system is stable for large enough *γ*_*A*_, i.e. when the feedback gain is sufficiently small.

## IV. Discussion

We derived a mechanistic model for a cellular population controller, which is the most common population control system in bacteria [25], [31]. We showed that, in this controller architecture, there is a trade-off between robustness to environmental disturbances and robustness to perturbations in available resources to the genetic circuit. Placing different genetic circuit components in multiple cells has been proposed as a possible solution to relieve some resource sharing effects for larger genetic circuits [15], [21], [32]. However, our analysis shows that when fluctuations in cellular resources are considered, it is not possible to tune the population controller to be both robust to extracellular disturbances and to intracellular disturbances. Thus, if distributed biomolecular systems are to be used to mitigate resource loading within a single cell in order to make larger genetic circuits, the population controller should be designed to ensure an acceptable level of robustness to environmental disturbances and to perturbations due to resource fluctuations.

## ACKNOWLEDGMENT

The authors thank Ted Grunberg for helpful discussions.

